# Danuglipron and orforglipron engage GLP-1R through materially different interfaces: an unbiased molecular dynamics study

**DOI:** 10.64898/2026.09.30.754652

**Authors:** Ahmet Oztekin, Sude Turker

## Abstract

GLP-1R is among the most heavily pursued targets in metabolic disease, but its natural agonist is a 30-residue peptide that must be injected. Two oral non-peptide agonists have reached late-stage trials. Cryo-EM studies have already shown that danuglipron and CHU-128, a close analogue of orforglipron, occupy the receptor differently, but not how either interface behaves over time. We ran 500 ns of unbiased all-atom molecular dynamics on GLP-1R bound to GLP-1, danuglipron (PF-06882961) and orforglipron (LY3502970), one trajectory each, in a POPC bilayer with the G protein removed. Although CHU-128 has been simulated before, its analogue, orforglipron, has not. Scored against contacts computed from the deposited structures, under a force field that shares no parameters with previous oral agonist studies, we recovered 80%, 86% and 79% above 50% occupancy. Indexing contact by chemical group, rather than by receptor residue, and normalising by molecular size, separates the two oral compounds: orforglipron carries ring systems dedicated almost entirely to one receptor domain, one of them directing 99.7% of its contact to the extracellular domain, whereas every group of danuglipron engages both. Danuglipron is correspondingly the outlier on measures that do not depend on a chosen threshold: only 33% of its contacts exceed 90% occupancy against 72% and 84%, and it is the only system whose occupancies remain unconverged at 100 ns. With single trajectories, these are observations rather than tested claims, but two drugs of the same class and comparable size occupy this receptor in structurally distinct ways.

## 1. Introduction

GLP-1 receptor agonists have changed how type 2 diabetes and obesity are treated [1–3]. GLP-1 itself is a 30-residue incretin that dipeptidyl peptidase-4 clears quickly, so the drugs in use are stabilised analogues given by injection, which limits adherence and distribution more than efficacy.

GLP-1R is a class B1 GPCR. Receptors in this family bind peptide agonists in two places: a large extracellular domain (ECD) holds the C-terminal part of the peptide, and the N-terminal part inserts into the transmembrane bundle and activates the receptor [4,5]. A small molecule cannot cover both sites, so how non-peptide agonists work at all is a real structural question. Cryo-electron microscopy has answered part of it, and has already shown that non-peptide agonists do not converge on one solution: danuglipron overlaps the GLP-1 site, whereas CHU-128 binds with a more open receptor conformation at the extracellular face [6], and orforglipron is held by the extracellular domain, extracellular loop 2, and the transmembrane helices together [7].

Danuglipron was discontinued in April 2025 following a case of possible drug-induced liver injury that resolved on withdrawal [8]; the finding has not been attributed to GLP-1R pharmacology, and nothing reported in this paper bears on it. Orforglipron remains in development; however, both compounds are informative for how non-peptide agonists are recognised at this receptor.

A structure tells you which residues lie within contact distance of a ligand, but not for how long. A contact that holds continuously and one that breaks and reforms two hundred times look identical in a structural figure, although they mean different things for residence time. Common wet lab techniques cannot resolve this per residue, but unbiased simulation can. We simulated all three agonists under identical conditions and compared their interfaces by size, by distribution across the receptor’s two domains, and by how long individual contacts survive. The informative comparison proved to be danuglipron versus orforglipron: those two are similar in size and of the same class, so any difference between them comes from molecular architecture.

## 2. Methods

### 2.1 Structures and preparation

Coordinates for the membrane came from the OPM database, which supplies membrane-aligned assemblies: 6×18 (GLP-1), 6×1A (danuglipron) and 6XOX (orforglipron) [9]. We kept only the receptor and its agonist.

We deleted the Gs heterotrimer and the crystallisation chaperones. The Gαs α5 helix sits in the receptor core and holds the transmembrane bundle active, whichever agonist occupies the orthosteric site, so keeping it would have imposed the same constraint on all three systems in the region we wanted to compare. Removal also made the depositions comparable, since 6XOX uses a Gs/Gi chimera with scFv16 and an unresolved α-helical domain and is not the same construct as the other two.

Unresolved loops were rebuilt with pdbfixer 1.12.0 [10], modelled atoms flagged in the B-factor column, and chain termini capped. Disulfides were assigned from deposited connectivity rather than by distance, with the build set to fail if any expected disulfide did not form. Hydrogens were placed with Reduce through pdb4amber rather than by tleap [11].

### 2.2 Ligand parameterisation and system assembly

Danuglipron (UK4, 41 heavy atoms) and orforglipron (V6G, 65 heavy atoms) were given GAFF2 atom types and AM1-BCC charges with antechamber from AmberTools 24.8 [12–15]; the semiempirical step was run in sqm under AM1. parmchk2 found no missing parameters for either, so nothing was estimated by analogy.

Each complex was embedded in a POPC bilayer with packmol-memgen and PACKMOL [16,17], solvated, and neutralised with 0.15 M KCl. We used ff19SB, Lipid21, GAFF2, and the OPC four-point water model against which ff19SB was parameterised [18–20]; topologies were converted to GROMACS format with ParmEd 4.3.1 [21]. The systems came to 139,445–139,900 atoms, 228–252 lipids and roughly 25,000 waters, in boxes near 100 × 100 × 140 Å.

### 2.3 Simulation protocol

Simulations were run in GROMACS 2026.3 (mixed precision, CUDA) [22]. Electrostatics used particle-mesh Ewald [23] with a 1.0 nm real-space cutoff and 0.12 nm grid spacing; van der Waals used a 1.0 nm cutoff with a potential-shift modifier and long-range dispersion correction.

Hydrogen mass repartitioning (3.024 Da, water excluded) allowed a 4 fs step [24], with bonds to hydrogen constrained by LINCS [25] at order 6 and 2 iterations rather than the usual 4 and 1. At 4 fs, the methionine S-methyl groups become the limiting motion. Default settings gave about 375 constraint warnings per nanosecond and terminated the runs, and the tighter setting removed them at roughly 4% throughput cost.

Temperature was held at 310.15 K by stochastic velocity rescaling [26] applied separately to solute, membrane, and solvent, since coupling them together equilibrates a bilayer to the wrong temperature. Pressure was held at 1 bar with semi-isotropic coupling. Every system went through the same staging: minimisation; 250 ps NVT with restraints; 2 ns restrained NPT under stochastic cell rescaling [27]; 10 ns unrestrained NPT while the bilayer relaxes; then 500 ns of production under Parrinello–Rahman [28].

### 2.4 Analysis

We discarded the first 50 ns of each run, allowing the receptor to relax from its cryo-EM starting conformation into the bilayer; everything reported comes from the 50–500 ns window, 4,501 frames at the trajectory’s 100 ps spacing. A contact is a receptor residue with any heavy atom within 4.0 Å of any ligand heavy atom, and occupancy is the percentage of analysed frames where that holds. Hydrogen bonds used 3.5 Å and 120°, salt bridges 4.0 Å. We used MDAnalysis 2.10.0 [29,30]. Trajectories were written for solute and membrane only, so water-mediated contacts could not be evaluated.

Contact tables indexed by receptor residue do not map onto the units a medicinal chemist works with, so we also split each agonist into chemical groups from its heavy-atom bond graph. Fundamental cycles were found from a spanning tree, and cycles sharing two or more atoms merged into fused ring systems. Ring systems joined by a single bond were kept separate, since that bond is the one a chemist would cut. Remaining acyclic atoms were grouped by connectivity, and per-group occupancy and domain distribution were computed as above.

Occupancy is estimated from a temporally correlated series, so its standard error depends on the correlation time of the contact indicator rather than the frame count. For each contact, we computed the integrated autocorrelation time τc of its binary indicator by the initial-positive-sequence estimator, and report σ = √[2 τc p(1 − p) / T]. Checked against the observed 100 ns to 500 ns change across the 97 contacts above 20% final occupancy, σ orders the contacts well but understates the size of the change: Spearman ρ is 0.886, the median observed-to-predicted ratio is 2.41, and 41% of contacts fall within 2σ. The understatement is expected, because the 100 ns to 500 ns change carries slow drift as well as statistical scatter, and a stationary autocorrelation estimate cannot capture the former. We therefore use σ to rank which occupancies to distrust rather than as an absolute error bar. Standard tools report occupancies as point values with no uncertainty; we report both.

For the structure–activity comparison in Section 3.8, we fixed a banding in advance: at a group held above 95% occupancy, a modification counts as a large loss at 10-fold or more, a moderate loss between 2- and 10-fold, and no loss below 2-fold. Potencies reported only as a bound are treated as qualitative. The thresholds are a stated convention, not a result.

The pipeline is scripted end-to-end, and no web-based system builder was used at any stage.

Development of the pipeline was assisted by a large language model (Claude, Anthropic), used to write and debug the build and analysis scripts, and generate the figures. All code was reviewed and executed by the authors, and every number reported here came from running that code on the trajectories; no value or citation was model-generated. The published potency data underlying Table 4 were curated by S.T. from the cited primary sources.

## 3. Results

### 3.1 Simulation quality

All three systems completed 500 ns without intervention (Figure 1, Table 1), and no agonist dissociated. Agonist RMSD was lowest for orforglipron at 1.59 Å. Receptor backbone RMSD had not plateaued in any system and was highest for 6XOX at 3.38 Å, which is expected for a ∼400-residue receptor relaxing out of a cryo-EM conformation, but means we restrict structural claims to the ligand pose. Bilayers settled near 71 Å^2^ per lipid, computed without subtracting the receptor cross-section and used only as an equilibration diagnostic.

**Table 1.**
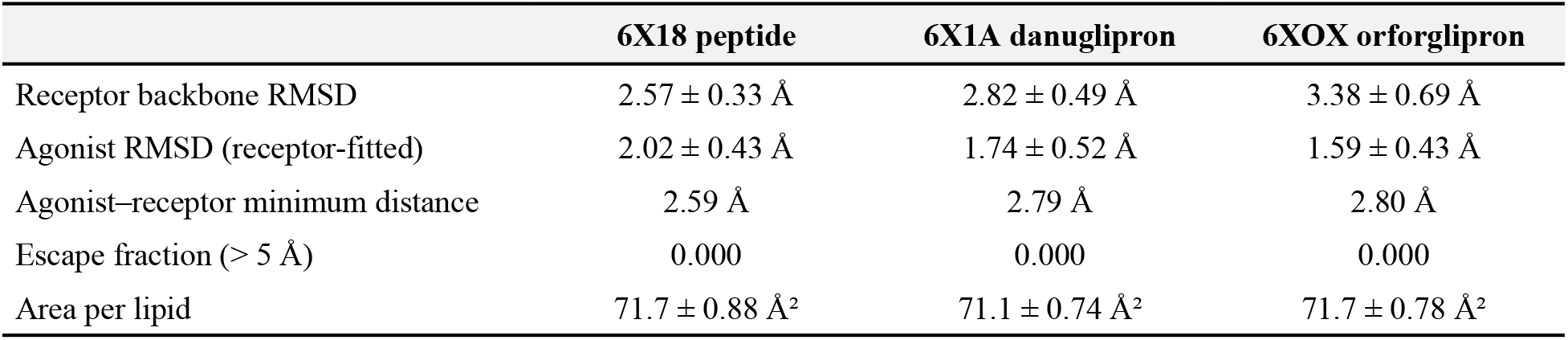
Simulation quality metrics over the 20–500 ns window (4,801 frames per system). This table reports equilibration quality and therefore uses a shorter discard than the 50 ns applied to every contact analysis below.

**Table 2.** Interface properties and convergence over the 50–500 ns window. Drift rows are computed over contacts above 20% final occupancy (n = 49, 24, and 24).

|  | Peptide | Danuglipron | Orforglipron |
| --- | --- | --- | --- |
| Residues contacted (> 50%) | 39 | 21 | 19 |
| — extracellular domain (24–145) | 17 | 6 | 7 |
| — transmembrane domain (146–423) | 22 | 15 | 12 |
| Contacts above 90% occupancy | 72% | 33% | 84% |
| Contacts above 99% occupancy | 38% | 19% | 37% |
| Hydrogen bonds above 50% occupancy | 13 | 2 | 1 |
| Salt bridges above 50% occupancy | 8 | 2 | 1 |
| Polar interactions per contacted residue | 0.54 | 0.19 | 0.11 |
| Median occupancy drift, 100 → 500 ns | −0.1 pts | −5.9 pts | +0.1 pts |
| Mean occupancy drift | −0.9 pts | −9.8 pts | +0.0 pts |
| Contacts overstated at 100 ns (< −5) | 12 | 13 | 3 |
| Contacts stable at 100 ns (± 5) | 32 | 10 | 17 |

**Figure 1.**
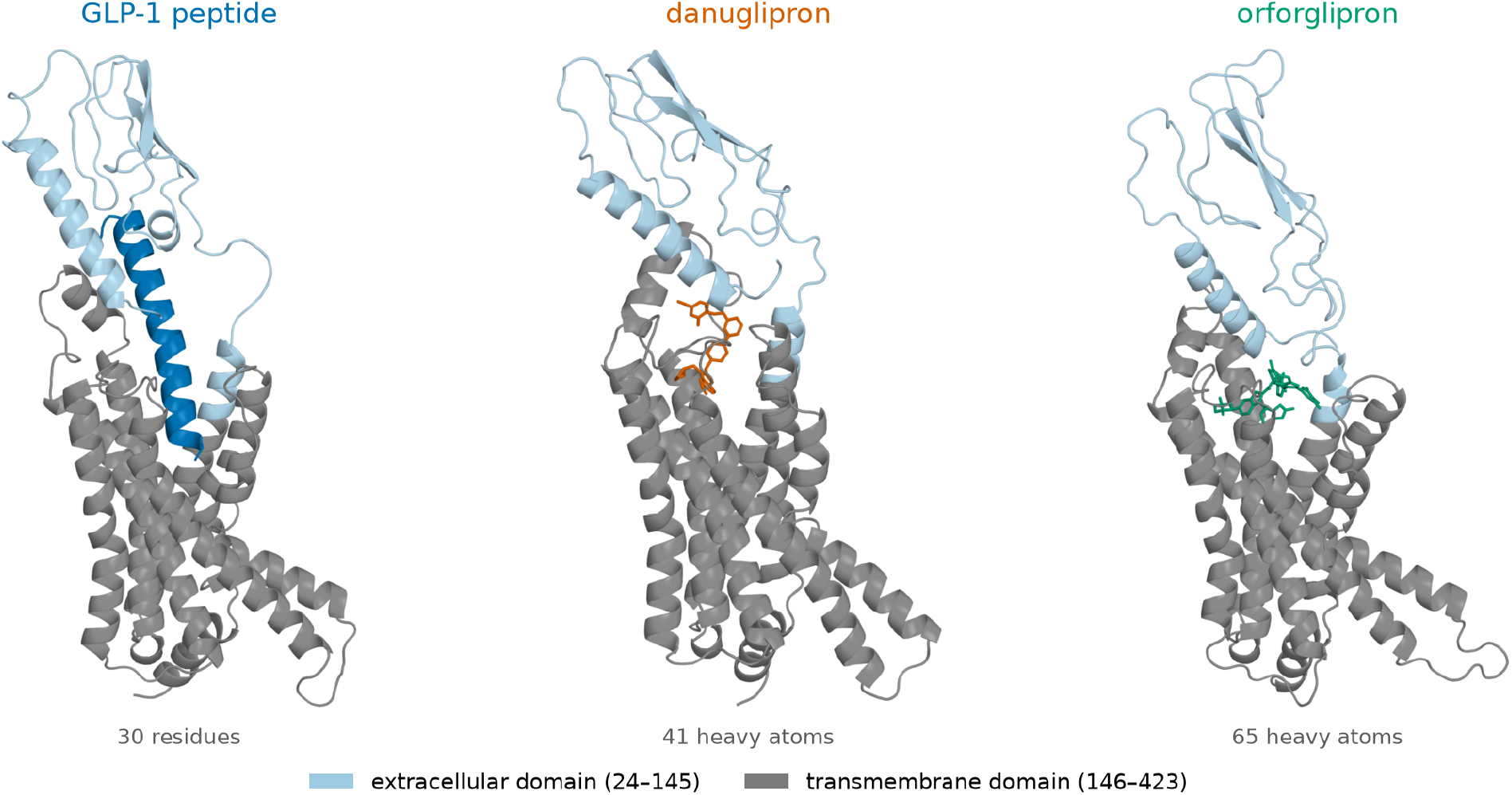
The three agonists in GLP-1R. Representative frames at 275 ns, receptor-superposed and shown in the OPM membrane frame with the extracellular domain uppermost. Receptor cartoon coloured by domain: extracellular (residues 24–145) light blue, transmembrane (146–423) grey. GLP-1 peptide as a dark blue cartoon; danuglipron and orforglipron as sticks.

**Figure 2.**
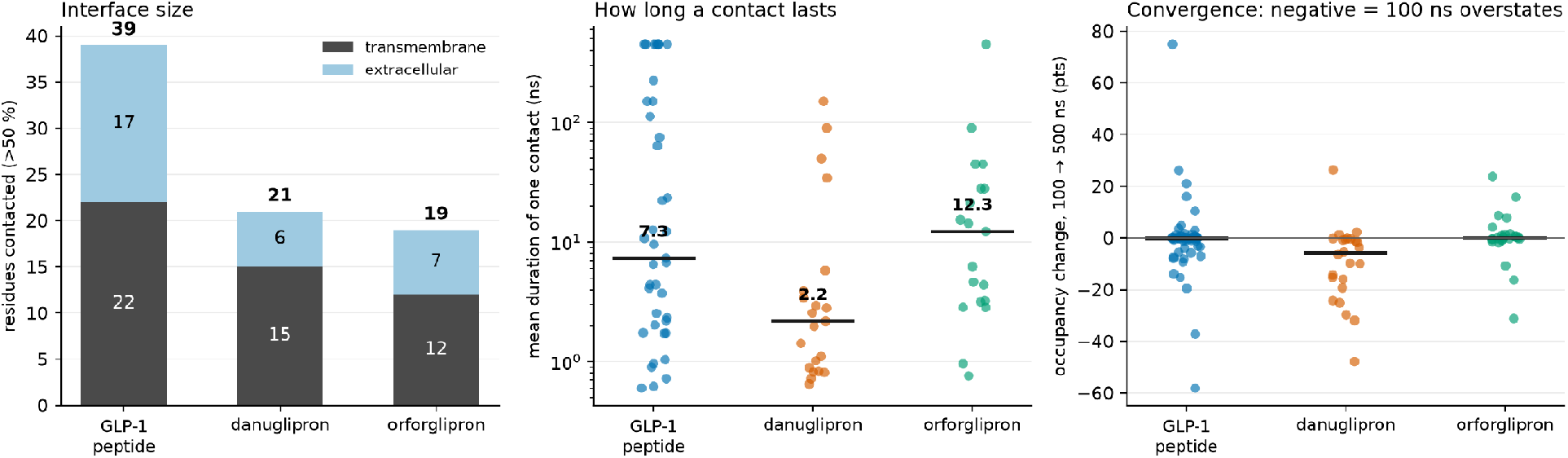
Interface size, contact-lifetime distributions and convergence. Left: residues contacted above 50% occupancy, partitioned by receptor domain. Centre: mean duration of an individual contact, one point per residue, horizontal bar at the median (logarithmic scale); note the dispersion, which the medians quoted in the text do not convey. Right: change in occupancy between a 100 ns and a 500 ns analysis window, one point per contact.

**Figure 3.**
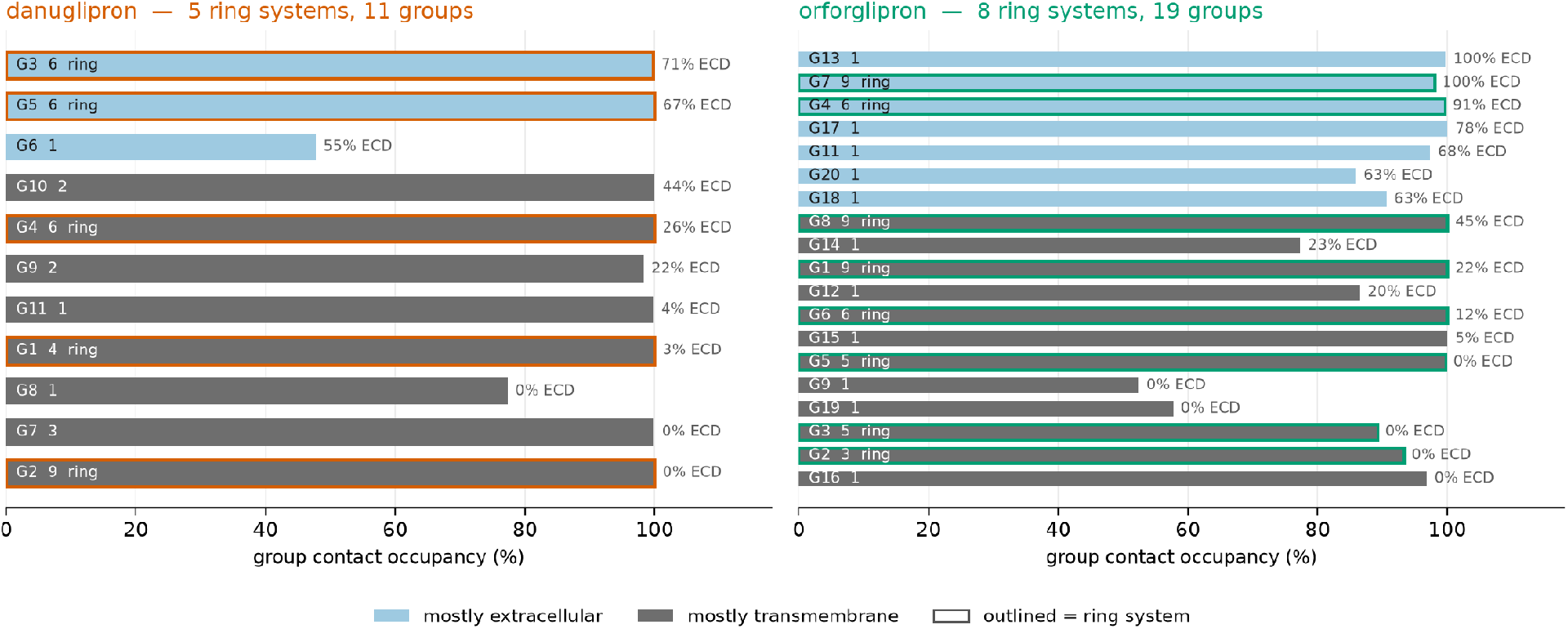
Ligand group decomposition. Each bar is one chemical group, either a fused ring system or an acyclic fragment, with its contact occupancy. Groups are ordered by the proportion of their contact directed at the extracellular domain; ring systems are outlined.

### 3.2 Recovery of experimental contacts

We computed contacts from the deposited coordinates under the same 4.0 Å criterion, giving 40 contacts for the peptide, 21 for danuglipron, and 19 for orforglipron. Above 50% occupancy, we recovered 32 (80%), 18 (86%) and 15 (79%); above 20%, 37 (93%), 20 (95%) and 17 (89%), respectively. Missed contacts are otherwise peripheral, the weakest being Val194 at 6% and Asp372 at 2%, with one complete miss: Glu34 is a deposited orforglipron contact that our trajectory never forms.

This is a positive control, not a finding, but it is worth something: the deposited structures were interpreted under CHARMM-family parameters while we used ff19SB, GAFF2 and OPC, so the agreement is force-field orthogonal, and no information about the reported contacts entered our simulations.

One result bears on how contact data should be compared with functional data. Alanine scanning of ECL2 identifies residues whose substitution shifts GLP-1 signalling toward pERK1/2 [31], and our peptide trajectory splits that set in two. Cys296, Trp297, Asn302, Tyr305 and Leu307 show zero occupancy and make no contact in the deposited 6×18 structure either, so their effect cannot be mediated by direct engagement of the peptide. Arg299 and Asn300, from the same scan, are contacted in essentially every frame at 100% and 99.3%, and Trp306, the one residue whose substitution abolished all biological activity, is contacted at 90.3%. Functional importance and direct contact are therefore neither equivalent nor mutually exclusive, and a set that conflates them cannot be scored against occupancy in either direction.

### 3.3 Interface composition

Interface size does not order the three agonists by molecular size. The peptide contacts roughly twice as many residues as either small molecule, but orforglipron is 59% larger than danuglipron by heavy-atom count (65 against 41) and contacts fewer residues, not more.

Composition separates these two oral agonists more than extent does. Only a third of danuglipron’s contacts exceed 90% occupancy against roughly three-quarters for the other two, so its interface is weighted toward intermediate occupancies. That is a property of its own distribution, not of a threshold we imposed.

Polar character orders them differently again, and inversely to persistence. The peptide maintains 13 hydrogen bonds and 8 salt bridges above 50% occupancy, several effectively permanent. Per residue contacted, that is 0.54 polar interactions, against 0.19 for danuglipron and 0.11 for orforglipron. Orforglipron therefore sustains the most uniformly high-occupancy interface of the three while forming the fewest polar contacts per residue engaged.

The two oral agonists engage substantially different residues; their contacts above 95% occupancy overlap at only two positions, Lys197 and Trp33. Orforglipron draws heavily on Leu141, Tyr145 and Glu138, at 99.6%, 99.3% and 98.7%. Of the three, only Tyr145 is absent from danuglipron’s interface altogether; Leu141 and Glu138 are held at 71.3% and 50.2%, so for those two the difference is one of degree rather than of identity. Trp33 is engaged at 99.8% by danuglipron and 96% by orforglipron. Domain-swap mutagenesis shows that Trp33 is required for orforglipron activity. Since the compound is inactive at the mouse receptor, mouse S33W restores activation and human W33S abolishes function [7]. Finding that residue among the most persistently engaged contacts is independent support for the simulation.

Wu and colleagues partition the small-molecule agonists structurally, and place danuglipron and orforglipron in different chemotypes engaging different helices: chemotype 3 interacting specifically with TM7 at Arg380, Phe381 and Leu384, chemotype 4 with TM1 at Pro137, Glu138, Leu144, Tyr145 and Tyr148 [38]. Our occupancies recover that partition at all eight residues. Danuglipron holds the three TM7 positions at 74.5%, 94.3%, and 89.4%, while orforglipron reaches none of them above 12%. Orforglipron holds the five TM1 positions at 98.2%, 98.7%, 37.8%, 99.3%, and 64.4%, while danuglipron contacts only Glu138 at 50.2%. The partition was derived from cryo-EM and druggability analysis rather than from dynamics, so its recovery here is independent.

Raising the threshold separates the two small molecules further. Orforglipron’s extracellular share barely moves, at 37%, 35% and 38% of contacts above 50%, 70% and 90% occupancy, while danuglipron’s runs 29%, 31% and then 14%.

### 3.4 Contact lifetimes

We report contact lifetimes separately because, unlike the compositional measures above, they are conditioned on where the occupancy threshold is placed, and the three ligands have different occupancy distributions, so a fixed cut samples a different part of each.

Median per-residue mean lifetime at a 50% threshold is 7.3 ns for the peptide, 2.2 ns for danuglipron, and 12.3 ns for orforglipron. At 70% it is 10.8, 2.9, and 14.3 ns, respectively; danuglipron is the shortest at both. At 90% the ordering breaks down, since its seven surviving contacts are its strongest and give the longest median of the three, and no pairwise difference is significant there (Mann–Whitney p = 0.28 to 0.74). We therefore make no claim that any one agonist forms longer-lived contacts than another.

Two cautions apply. Seven peptide contacts and one orforglipron contact stay unbroken across the whole 450 ns window, so their lifetimes are right-censored lower bounds; excluding them lowers the 50% median from 7.3 to 4.4 ns for the peptide and 12.3 to 9.2 ns for orforglipron, leaving danuglipron unchanged. Interquartile ranges are also very wide, spanning nearly two orders of magnitude for the peptide.

### 3.5 Ligand group decomposition

Every ring system in both molecules is engaged for the great majority of the trajectory, the least-engaged at 99.8% occupancy for danuglipron and 89.2% for orforglipron, so neither carries a structurally passive scaffold. Raw counts scale with size and carry no independent information: ring density is 0.122 and 0.123 per heavy atom, and 76% and 80% of heavy atoms lie within rings. Orforglipron is larger because it has more rings.

How those groups distribute between the domains is another matter. Normalised by heavy-atom count, orforglipron directs 0.108 groups per heavy atom predominantly at the extracellular domain, against danuglipron’s 0.073. More importantly, the two differ categorically in whether any group is domain-specific at all: orforglipron’s G7 directs 99.7% of its contact to the extracellular domain, and G4 directs 90.9%, while G2 and G3 are entirely transmembrane. Danuglipron differs from the other two: its most extracellularly directed ring still spends 29% of its contact inside the transmembrane domain.

That fits the threshold behaviour stated before: for orforglipron, extracellular engagement is carried by groups dedicated to that domain and survives into its strongest contacts, while danuglipron’s is spread across groups that also engage the transmembrane bundle.

Clustering of per-frame contact fingerprints resolves no distinct binding modes in any system (optimal k = 2 throughout, silhouette 0.071–0.129), so contact-event counts reflect fluctuation within a single pose rather than exchange between poses.

### 3.6 Dependence of occupancy on trajectory length

Recomputing every occupancy over a growing window shows the three systems converge differently (Figure 4). Danuglipron moves most, and unlike the lifetime comparison, this ordering is stable under the choice of occupancy floor: median drift is −5.9, −4.6, and −4.6 percentage points for contacts above 20%, 40%, and 60% final occupancy, against values within 0.1 points of zero for the other two. Glu138 registers 98.0% at 100 ns and 50.2% at 500 ns.

**Figure 4.**
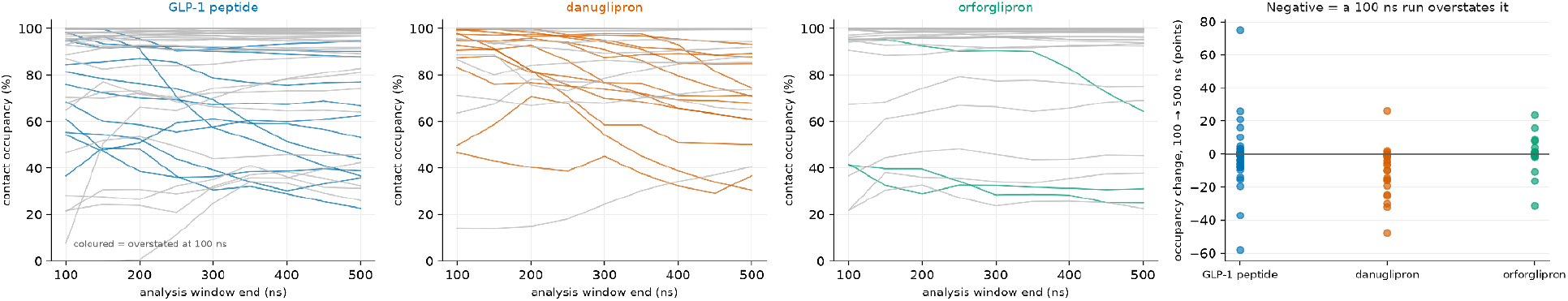
Occupancy as a function of analysis-window length. Each line is one contact, recomputed over analysis windows ending between 100 and 500 ns. Coloured lines denote contacts overstated by more than 5 percentage points at 100 ns.

This follows from the kinetics: a contact that never ruptures cannot drift, whereas one that reforms 175 times needs many events before its occupancy settles. We therefore treat our own occupancies for short-lived interfaces as provisional unless a convergence check accompanies them, and the same caution seems worth applying to other non-peptide ligands.

### 3.7 Contacts absent from the deposited structures

Seven residues are engaged above 50% occupancy in our trajectories yet fall outside the deposited contact set of that ligand (Table 3). These are what the dynamics add to the static picture, and they are directly testable.

**Table 3.** High-occupancy contacts absent from the corresponding deposited structure.

| Agonist | Residue | Occupancy | Mean lifetime | Events | Note |
| --- | --- | --- | --- | --- | --- |
| Orforglipron | Thr298 | 99.3% | 27.95 ns | 16 | deposited contact for danuglipron, not orforglipron |
| Orforglipron | Asp198 | 95.6% | 4.63 ns | 93 | — |
| Orforglipron | Ala200 | 95.3% | 4.38 ns | 98 | ECL1 hydrophobic network with Tyr220 |
| Orforglipron | Leu142 | 69.2% | 0.76 ns | 412 | — |
| Danuglipron | Arg299 | 88.1% | 2.81 ns | 141 | ECL2; R299A alters signalling bias |
| Danuglipron | Leu218 | 68.0% | 0.72 ns | 424 | — |
| Danuglipron | Glu138 | 50.2% | 1.02 ns | 221 | deposited contact for orforglipron, not danuglipron |

**Table 4.** Published potencies for analogues varying orforglipron group G8, joined to the occupancy of that group. EC50 by cAMP accumulation [37]; fold is against (R/S)-orforglipron at 1.1 nM. Class follows the banding in Section 2.4, and (b) marks a bound rather than a measurement. Listed are 24 of 32 analogues that lose 2-fold or more

| Cpd | EC50 (nM) | Fold | Class | Cpd | EC50 (nM) | Fold | Class |
| --- | --- | --- | --- | --- | --- | --- | --- |
| 1 | >100 | >91 | ≥10× (b) | 16 | 12.0 | 10.9 | ≥10× |
| 2 | >100 | >91 | ≥10× (b) | 18 | 6.0 | 5.5 | 2–10× |
| 5 | 25.0 | 22.7 | ≥10× | 19 | 20.0 | 18.2 | ≥10× |
| 6 | 2.7 | 2.5 | 2–10× | 20 | >100 | >91 | ≥10× (b) |
| 7 | 38.0 | 34.5 | $\geq 10\times$ | 21 | 5.6 | 5.1 | 2–10× |
| 8 | 3.3 | 3.0 | 2–10× | 23 | 12.0 | 10.9 | $\geq 10\times$ |
| 9 | 2.2 | 2.0 | 2–10× | 25 | 4.6 | 4.2 | 2–10× |
| 10 | 8.0 | 7.3 | 2–10× | 26 | 44.0 | 40.0 | $\geq 10\times$ |
| 11 | 3.4 | 3.1 | 2–10× | 27 | 21.0 | 19.1 | $\geq 10\times$ |
| 12 | 15.0 | 13.6 | $\geq 10\times$ | 28 | 2.6 | 2.4 | 2–10× |
| 13 | 39.0 | 35.5 | $\geq 10\times$ | 29 | 17.0 | 15.5 | $\geq 10\times$ |
| 15 | 4.0 | 3.6 | 2–10× | 32 | 4.2 | 3.8 | 2–10× |

Thr298 and Glu138 are the informative entries because they exchange between the two compounds: each is a deposited contact of one ligand and a sustained simulated contact of the other. The danuglipron structure lists Thr298, and the orforglipron structure does not; yet orforglipron holds it for 99.3% of the run and danuglipron for 15.7%. Glu138 runs the other way: it is listed for orforglipron, not for danuglipron, yet danuglipron reaches it at 50.2%. Thr298 is not new to the orforglipron complex; Kawai and colleagues describe its hydrogen-bonding to the Trp33 lid that closes over the indole-tetrahydropyran branch [7], but it is not within 4.0 Å of the ligand in the deposited coordinates, so what the trajectory adds is its promotion from a residue stabilising that lid to a direct and near-permanent ligand contact. Glu138 is the weaker of the two entries: at 50.2% it clears our threshold by 0.2 points, and it is the least converged occupancy we report (Section 3.6), so it should be read as provisional. Alanine substitution at these positions should therefore affect the two compounds asymmetrically.

Ala200 belongs to a hydrophobic network stabilising the TM2–ECL1–TM3 region that also includes Tyr220, orforglipron’s single most persistent contact at 100% occupancy across an unbroken 450 ns. Engaging both members of that pair suggests orforglipron’s grip on ECL1 is more extensive than the deposited structure indicates.

### 3.8 Structure–activity data at a maximally engaged group

A published series lets us ask what modifying one maximally engaged group costs in potency. Li and colleagues report 32 analogues of orforglipron varying a single branch, assayed by cAMP accumulation [37]. Because the (S)-configured starting material was initially unavailable, compounds 1–32 were made from racemic material and tested as epimeric mixtures, so the authors compare against (R/S)-orforglipron at 1.1 nM, which we use as the reference throughout. The same paper reports authentic orforglipron at 0.51 nM, against which every fold change below would roughly double. Once the (S) intermediate became available, the authors resynthesised their three most potent compounds as single epimers, which fall outside the 32 analysed here.

The branch varied is the 4-fluoro-1-methylindazole, which our decomposition assigns to group G8. Li and colleagues describe this branch as solvent-exposed [37]; the two descriptions are compatible, since occupancy counts contact rather than burial, and a group can be the most solvent-accessible part of a ligand while still holding receptor contact in every frame. G8 is engaged in every analysed frame and holds Lys202 in 98% and Glu138 in 96%. Its contacts fall in both domains, so this series does not bear on the domain-specialisation argument in either direction.

Across the full series, 13 analogues lose 10-fold or more and 11 lose between 2- and 10-fold (Table 4). The remaining eight fall within 2-fold of the parent, and compound 14 is more potent at roughly 0.4 nM. One point does not come from us: Li and colleagues attribute the inactivity of two analogues to Lys202 [37], the residue our decomposition independently shows G8 gripping hardest.

Two limits apply. Every compound varies the same group, so the series characterises one group in depth and cannot test whether occupancy predicts potency across groups. And because a quarter of the series is unaffected by modification there, the consequence of altering a group held in essentially every frame ranges from nothing to more than 90-fold. Occupancy identifies where a ligand leaves no spare room, but it does not say what any contact in that region is worth.

Danuglipron has no comparable dataset. The only series varying its groups one at a time reaches us third-hand, as fold shifts read from a figure in a preprint [38] that took them from patent WO2018109607A1, with no EC50 value and no assay stated for any of its three rows. Taken at face value, they agree with the orforglipron picture, but third-hand shifts without a stated assay carry little weight, and we report those rows in the supporting data rather than as a table. The main medicinal-chemistry series cannot be used at all [39], since every step changes more than one group at once or crosses assay formats.

What neither ligand supplies is a negative control. Every group with any structure–activity coverage sits above 89% occupancy, so only half our stated convention is ever exercised. These data can show that modifying a maximally engaged group sometimes costs a great deal; however, they cannot show that modifying a weakly engaged group costs less, which is the comparison that would make occupancy predictive rather than descriptive.

## 4. Discussion

The comparison that carries most information is between the two non-peptide agonists. Danuglipron and orforglipron have comparable mass, the same target, and the same therapeutic class, yet occupy overlapping but substantially distinct residue sets, and their interfaces differ systematically in composition. Danuglipron is the outlier: only a third of its contacts exceed 90% occupancy where roughly three-quarters of the other two do, its extracellular engagement does not survive at high occupancy where orforglipron’s does, and it is the only system substantially unconverged at 100 ns.

That result has independent support. Xu and colleagues simulated the same complex under CHARMM36m and report notable conformational deviation of the ligand after 200 ns across three independent replicates [32]. Under a force field sharing no parameters with Xu and colleagues’ simulation, and from a single trajectory, we find the least converged interface of the three. This comparison is not unprecedented in its chemotype. Xu and colleagues pair danuglipron with CHU-128 and Peptide 19 [32], and Brubaker and colleagues pair it with CHU-128, semaglutide, and tirzepatide [33]. CHU-128 is a close analogue of orforglipron: both carry 65 heavy atoms and the same pyrazolo[4,3-c]pyridine core, indazole, and oxadiazolone, differing at the indole substituent and the linker. Wu and colleagues place the two in one chemotype and compare their bound poses moiety by moiety [38]. Those studies therefore already set danuglipron against this chemotype, and both found danuglipron to be the less stable of the pair, as Zhang and colleagues had found it to be the more GLP-1-like [6]. What has not been done is to simulate the clinical molecule itself: neither study includes orforglipron, for which MemProtMD holds only an automated pipeline assembly at coarse-grained and short atomistic resolution, built for lipid-interaction annotation rather than agonist comparison [34]. That three independent force fields and one cryo-EM series separate these two compounds in the same direction is better evidence than any one of them alone.

The orforglipron series adds a second check, from chemistry rather than structure: the residue our decomposition shows the indazole branch gripping hardest, Lys202, is the residue Li and colleagues invoke to explain why two analogues are inactive [37]. What that series does not do is tell us what occupancy is worth, since every compound in it varies a group held in essentially every frame and the consequence still ranges from nothing to more than 90-fold.

Interface persistence does not follow the properties we expected. Orforglipron sustains the most uniformly high-occupancy interface while forming the fewest polar interactions per residue engaged, and contacts fewer residues than danuglipron despite being 59% larger. Neither size nor polar anchoring orders the three agonists.

What distinguishes them, once size is divided out, is how ligand substructure is distributed between the receptor’s two domains. Our reading is that a ligand anchored at points in structurally independent regions is released only when those points release together, whereas one whose contacts all lie within a single region stays vulnerable to a local fluctuation that interrupts several at once. That phenomenon shows up in the threshold behaviour of the extracellular share and in the convergence behaviour, and both separate the two compounds in the same direction.

Two implications follow, if the pattern holds. For design, what appears to matter is where ligand substructure sits rather than how strongly any single contact forms, which argues for spreading contact across both domains rather than strengthening the transmembrane pocket. For mechanism, it suggests how a small molecule partly recapitulates the two-domain engagement the peptide achieves by spanning the receptor: by placing distinct substructures in each domain instead of reaching across.

Indexing occupancy by chemical group rather than by receptor residue is what made this visible, and normalising by molecular size keeps it from restating molecular weight. The same holds for the uncertainties we attach to every occupancy: most tools emit point values with no error estimate, and here discrepancies above 40 percentage points separate 100 ns and 500 ns windows for short-lived contacts.

### 4.1 Limitations

These results come from a single trajectory per system. We describe the differences in interface size, persistence, and convergence; however, we have not tested them. With no replicate-level variance, these claims about the MD runs should be read as observations about these trajectories, not as supported claims about the molecules. Contact frequencies vary between independent replicas of the same system [35,36], and the size of what we report has to be weighed against that. A replicate campaign is underway.

Receptor backbone RMSD had not plateaued in any system and was highest for orforglipron at 3.38 Å, so we can make no claims about receptor conformation. We cannot exclude that orforglipron’s distinctive interface reflects a receptor still relaxing around it, though it has the lowest agonist RMSD and the best-converged occupancies, neither expected of an interface still reorganising.

We removed the G protein by design, so no result here bears on receptor activation, transducer coupling, signalling bias or efficacy. Contact occupancy is not a binding free energy, and nothing in this work predicts potency.

Because production trajectories excluded solvent, we could not examine water-mediated contacts. Zhang and colleagues show these form extensive networks below the PF-06882961 site and discriminate between GLP-1R agonists [6].

The contacts in Table 3 are predictions. They may reflect genuine sampling beyond the cryo-EM conformation, force-field bias, or incomplete relaxation, and have not been tested experimentally.

### 4.2 Outlook

Replication is the immediate priority, and independent trajectories will turn these comparisons from descriptive into statistical ones.

After that, the exchange of Thr298 and Glu138 between the two compounds (Table 3) is the most specific test available: each is a reported contact of one ligand and a sustained contact of the other, so alanine substitution should affect them asymmetrically, with T298A reducing orforglipron potency more than the deposited structures predict and E138A affecting danuglipron. The T298A arm is the stronger of the two, since Glu138’s occupancy in danuglipron is both marginal and the least converged we report, and a replicate campaign should settle whether it survives. Because the prediction is directional and concerns two compounds, it is not easily satisfied by chance.

## Supporting information

SAR Table

## Author contributions

**A.O**.: conceptualisation, methodology, software, investigation (molecular dynamics simulations), formal analysis, visualisation, writing.

**S.T**.: methodology (design of the structure–activity table and its inclusion criteria), data curation (extraction of published potency data, source provenance assessment, and compound-to-group mapping), formal analysis, validation, writing, review and editing.

All authors have read and agreed to the published version of the manuscript.

## Data availability

All materials behind this work are available from the corresponding author on reasonable request: the build and analysis scripts, the input structures, the ligand parameters and mdp files, every analysis output behind the figures and tables, the production trajectories, and the curated structure–activity dataset underlying Table 4 with its source provenance and compound-to-group mapping.

## Acknowledgements

The authors used Claude (Anthropic) for assistance with code development and manuscript drafting. The authors reviewed and edited all output and take full responsibility for the content of this work.

