## Supplementary material for "Danuglipron and orforglipron engage GLP-1R through materially different interfaces: an unbiased molecular dynamics study": SAR Table

| Compound | Modification (Ar group) | EC50 (nM) | Fold | Assay | Group | Occ. | Holds | Consistent | Source |
|---|---|---|---|---|---|---|---|---|---|
| orforglipron | Parent. 4-fluoro-1-methylindazol-5-yl | 0,51 | 1,00 | cAMP | — | | — | — | Li et al. 2026, Table 1 |
| (R/S)-orforglipron | Same Ar; epimeric pair at the tetrahydropyran 4-position | 1,10 | 1,00 | cAMP | — | | — | — | Li et al. 2026, Table 1 |
| 1 | Benzimidazole, linearly fused to a 5-membered ring | >100 | | cAMP | G8 — indazole branch | %100,0 | Lys202 98%, Glu138 96% | Yes (qual.) | Li et al. 2026, Table 1 |
| 2 | Benzimidazole, linearly fused to a 6-membered ring | >100 | | cAMP | G8 — indazole branch | %100,0 | Lys202 98%, Glu138 96% | Yes (qual.) | Li et al. 2026, Table 1 |
| 5 | Benzimidazole, linearly fused to a 7-membered ring | 25,00 | 22,73 | cAMP | G8 — indazole branch | %100,0 | Lys202 98%, Glu138 96% | Yes | Li et al. 2026, Table 1 |
| 6 | Benzimidazole, bent fused to a 6-membered aromatic ring | 2,70 | 2,45 | cAMP | G8 — indazole branch | %100,0 | Lys202 98%, Glu138 96% | Partial | Li et al. 2026, Table 1 |
| 7 | Dibenzofuran | 38,00 | 34,55 | cAMP | G8 — indazole branch | %100,0 | Lys202 98%, Glu138 96% | Yes | Li et al. 2026, Table 1 |
| 8 | Indoline-3-one, bent fused to 5-membered aromatic ring | 3,30 | 3,00 | cAMP | G8 — indazole branch | %100,0 | Lys202 98%, Glu138 96% | Partial | Li et al. 2026, Table 1 |
| 9 | Indazole-3-one, bent fused to 5-membered ring | 2,20 | 2,00 | cAMP | G8 — indazole branch | %100,0 | Lys202 98%, Glu138 96% | Partial | Li et al. 2026, Table 1 |
| 10 | N-methylindole, bent fused to 5-membered ring | 8,00 | 7,27 | cAMP | G8 — indazole branch | %100,0 | Lys202 98%, Glu138 96% | Partial | Li et al. 2026, Table 1 |
| 11 | N-methylindole, bent fused to 6-membered ring | 3,40 | 3,09 | cAMP | G8 — indazole branch | %100,0 | Lys202 98%, Glu138 96% | Partial | Li et al. 2026, Table 1 |
| 12 | Imidazo [1,2-a]pyridine, bent fused to cyclopentane | 15,00 | 13,64 | cAMP | G8 — indazole branch | %100,0 | Lys202 98%, Glu138 96% | Yes | Li et al. 2026, Table 1 |
| 13 | Indole, bent fused to 6-membered thiazine | 39,00 | 35,45 | cAMP | G8 — indazole branch | %100,0 | Lys202 98%, Glu138 96% | Yes | Li et al. 2026, Table 1 |
| 15 | Imidazole, bent fused to 6-membered N-methylpiperazine | 4,00 | 3,64 | cAMP | G8 — indazole branch | %100,0 | Lys202 98%, Glu138 96% | Partial | Li et al. 2026, Table 1 |
| 16 | Imidazole, bent fused to 6-membered 1,3-oxazine | 12,00 | 10,91 | cAMP | G8 — indazole branch | %100,0 | Lys202 98%, Glu138 96% | Yes | Li et al. 2026, Table 1 |
| 18 | 7-fluoro-benzimidazole fused to 1,4-oxazine | 6,00 | 5,45 | cAMP | G8 — indazole branch | %100,0 | Lys202 98%, Glu138 96% | Partial | Li et al. 2026, Table 1 |
| 19 | 7-fluoro-benzimidazole fused with Spiro[4,2] septane | 20,00 | 18,18 | cAMP | G8 — indazole branch | %100,0 | Lys202 98%, Glu138 96% | Yes | Li et al. 2026, Table 1 |
| 20 | 7-fluoro-benzimidazole fused to 2,6-dimethyl morpholine | >100 | | cAMP | G8 — indazole branch | %100,0 | Lys202 98%, Glu138 96% | Yes (qual.) | Li et al. 2026, Table 1 |
| 21 | Benzo-fused 1-methyl-1.4-diazepane | 5,60 | 5,09 | cAMP | G8 — indazole branch | %100,0 | Lys202 98%, Glu138 96% | Partial | Li et al. 2026, Table 2 |
| 23 | Benzo-fused methylazepane, inward N-methyl | 12,00 | 10,91 | cAMP | G8 — indazole branch | %100,0 | Lys202 98%, Glu138 96% | Yes | Li et al. 2026, Table 2 |
| 25 | Benzo-fused N-methyl-1,4-oxazepine | 4,60 | 4,18 | cAMP | G8 — indazole branch | %100,0 | Lys202 98%, Glu138 96% | Partial | Li et al. 2026, Table 2 |
| 26 | Benzo-fused N-methyl-1,4-oxazepan-5-one | 44,00 | 40,00 | cAMP | G8 — indazole branch | %100,0 | Lys202 98%, Glu138 96% | Yes | Li et al. 2026, Table 2 |
| 27 | Benzo-fused N-methyl-1,4,5-oxathiazepine | 21,00 | 19,09 | cAMP | G8 — indazole branch | %100,0 | Lys202 98%, Glu138 96% | Yes | Li et al. 2026, Table 2 |
| 28 | Benzo-fused N-ethyl-1,4-oxazepine | 2,60 | 2,36 | cAMP | G8 — indazole branch | %100,0 | Lys202 98%, Glu138 96% | Partial | Li et al. 2026, Table 2 |
| 29 | Benzo-fused N-cyclopropyl-1,4-oxazepine | 17,00 | 15,45 | cAMP | G8 — indazole branch | %100,0 | Lys202 98%, Glu138 96% | Yes | Li et al. 2026, Table 2 |
| 32 | Fluorobenzo-fused N-methyl-1,4-oxazepine | 4,20 | 3,82 | cAMP | G8 — indazole branch | %100,0 | Lys202 98%, Glu138 96% | Partial | Li et al. 2026, Table 2 |
